# Phototriggered proton-selective carrier currents through photoswitchable membranes

**DOI:** 10.1101/2025.01.13.632814

**Authors:** Juergen Pfeffermann, Rohit Yadav, Toma Glasnov, Oliver Thorn-Seshold, Peter Pohl

## Abstract

Controlling the flux of specific ions across lipid bilayers is a central challenge for cellular life. Here, we present a general strategy for the precise, noninvasive, and reversible photoregulation of specific ion transport across lipid bilayer membranes: using embedded azobenzene-containing photolipids as dopants that effectively photo-control the membrane permeation of specific ion carriers. Photolipid (OptoDArG) embedding and photoisomerization causes only minor changes in background ion conductance, consistent with classical models of membrane transport wherein small changes in bilayer thickness have little effect on ion permeability. Yet, in the presence of ion carriers, photoisomerization instead results in order-of-magnitude jumps in ion flux. These jumps are observed with both cationic and anionic carriers; particular protonophores deliver exceptional performance, with proton currents amplified by up to ≈200-fold upon UV illumination yet reversed by blue light exposure. A model in which interfacial uptake and release reactions are modulated by light-induced membrane changes explains these results. Doping photolipids together with well-established selective ion carriers into membranes thus presents a flexible and general chemical strategy for reversible, millisecond-scale photoregulation of specific ion transport across lipid bilayers. It achieves large currents, long-lasting effects, its ion scope is easily tuned by carrier choice, no genetic modifications are needed, and it has the potential to harness recent photoswitch advances to rationally shift its single-photon action spectra from the UV/Vis right up to the NIR/SWIR: a convenient all-chemical approach toward light-based ion permeability regulation in the biological or synthetic cell.

Graphical abstract

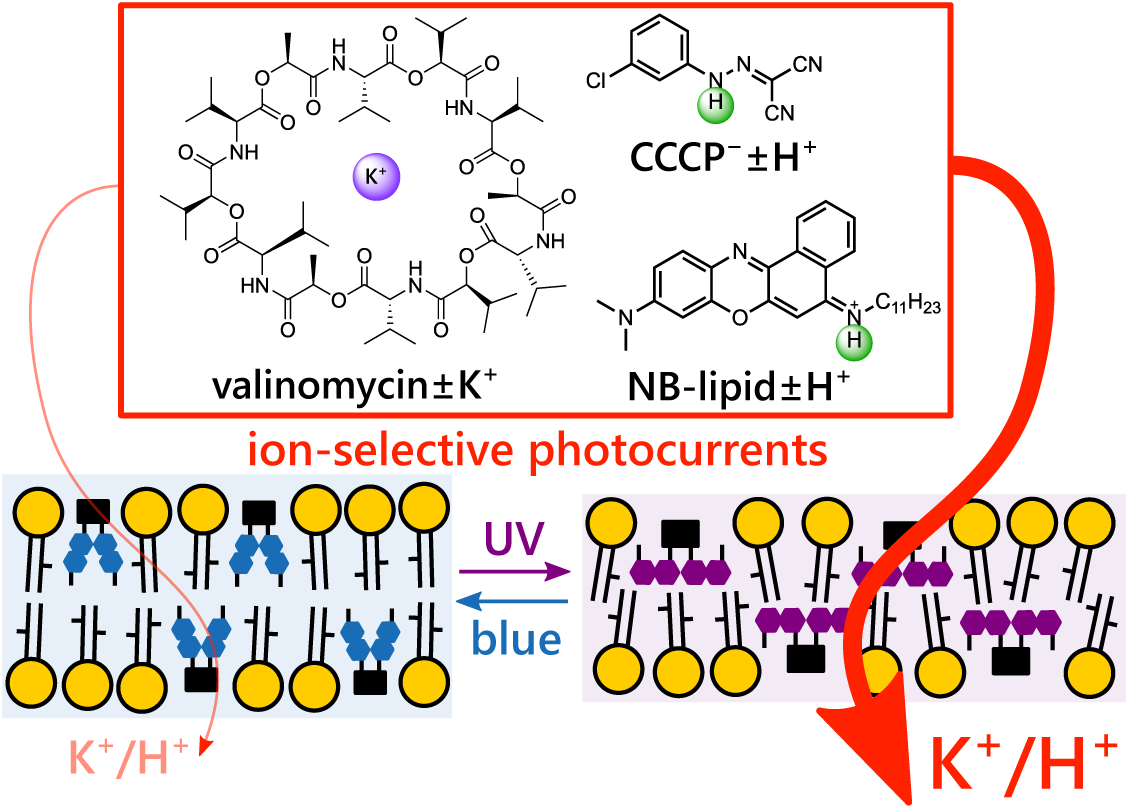

## Introduction

The regulated transport of ions across cellular and subcellular membranes is essential for life. For example, neuronal excitability depends on rapid changes in the permeability of sodium and potassium ions achieved through gating of ion-selective protein channels. In bioenergetics, proton permeability is adjusted by proteinaceous carriers to optimize ATP production while preventing the excessive generation of reactive oxygen species. Conceivably, the regulation of membrane ion transport by exogenous stimuli is of both scientific and pharmaceutical interest. To that effect, light has emerged as a desirable stimulus given the site-specific, minimally invasive, temporally precise, and reversible manner by which it may regulate membrane ion permeability, P. The latter is viewed as the proportionality factor between the transmembrane flux, J, and the transmembrane concentration gradient of an ion, Δc, in steady-state, whereby J = *−*P *·* Δc [1].

In a seminal article from 1969, Parsegian [2] outlined that the energetic barrier for an ion crossing the low-dielectric membrane cannot be effectively altered by changing membrane thickness or by ion pair formation. Instead, he recognized that mobile carriers and channels are effective in modulating P (Figure 1a) [2]. Channels provide a contiguous pathway of high polarizability through the membrane, containing water molecules and a polar interior. Mobile carriers, such as hydrophobic ionophores, facilitate the transport of ions across membranes by selective binding of ions, which lowers the electrostatic Born energy associated with the transfer of an ion into the interior of the hydrophobic membrane [3]; the basic carrier transport scheme is given in Figure 1a. Although Parsegian’s elegant concepts have withstood the tests of time, we report here that we can produce large alterations in P determined by selective ion carriers accompanying small light-induced changes in membrane thickness. Using several complementary experimental approaches, we seek to contour the underlying molecular determinants.

**Figure 1:**
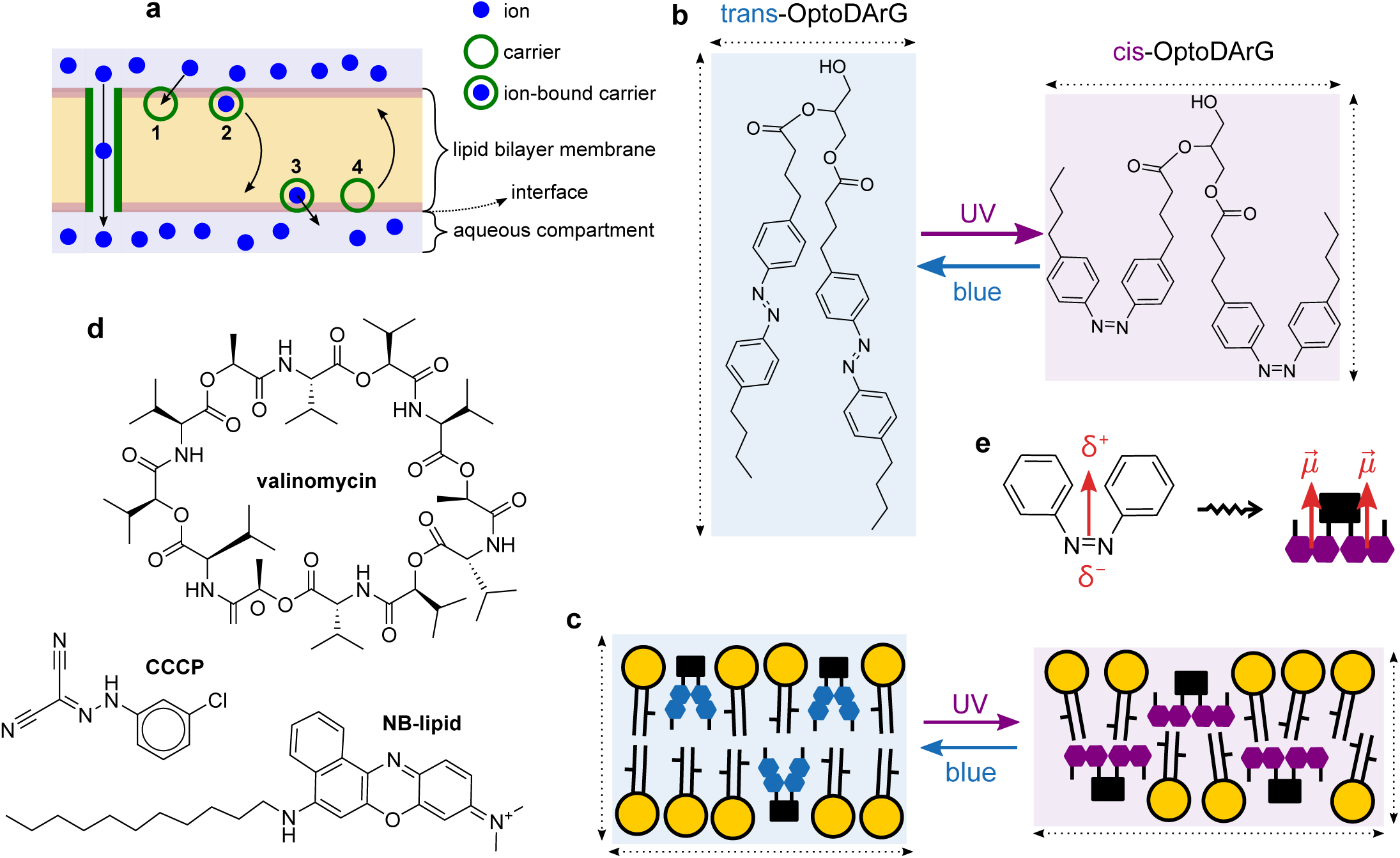
Components and concepts underlying photolipid-regulated carrier transport. **a,** Channels and small-molecule carriers facilitate transmembrane ion transport across biological membranes along the electrochemical gradient [2]. Carrier transport typically occurs in four steps: **1,** an ion at the bilayer–aqueous interface associates with an interfacially-adsorbed carrier molecule; **2,** the carrier– ion complex traverses the membrane; **3,** the ion dissociates from the carrier and is released into the membrane’s aqueous surroundings; **4,** the free carrier can traverse the membrane as well; it is free to bind another ion or recycle to the original side. **b,** The photolipid OptoDArG in its trans and cis state: blue light (488 nm) generates mainly the former, UV light (375 nm) generates mainly the latter. For brevity, the photoequilibria will henceforth be indicated as simply “trans” (blue light) and “cis” (UV light). cis-OptoDArG is broader and shorter than trans-OptoDArG. **c,** Both photoisomers incorporate into lipid bilayers and cellular membranes. Photoisomerization of membrane-embedded photolipids causes changes in global material properties owing to changes in molecular structure (cf. panel **b**). Structurally, photoisomerization to cis-OptoDArG by UV light increases bilayer surface area but reduces bilayer thickness. **d,** Chemical structures of the carriers used in this study: the cationic K^+^ ionophore valinomycin, the anionic protonophore CCCP and NB-lipid, a lipidated Nile Blue derivative which acts as a cationic protonophore. **e,** Cis-azobenzene has a dipole moment, 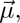 of magnitude 3 D pointing in the indicated direction; δ*^−^* and δ^+^ indicate negative and positive partial charge. A simplified view of the expected orientation of cis-OptoDArG in the lipid bilayer (cf. panel **c**) indicates that, on average, the azobenzene moieties’ dipole moments point towards the interfaces.

### Regulation of valinomycin–K^+^ flux across photoswitchable bilayers

Our approach leverages previous findings that the photolipid OptoDArG (Figure 1b) spontaneously incorporates into lipid bilayers and can undergo rapid (millisecond-scale) isomerization by exposure to intense UV (375 nm) or blue (488 nm) light there (irradiance of several hundred W cm*^−^*^2^). Consequent changes in molecular structure alter the material properties of the embedding bilayer membrane (Figure 1c) [4, 5]. We conducted initial experiments with the electrically neutral dodecadepsipetide macrocycle valinomycin (Figure 1d), an established K^+^-selective carrier [3, 6] synthesized by *Streptomyces fulvissimus* [7]. Valinomycin can permeate the membrane both without K^+^ and as a valinomycin–K^+^ complex, a hydrophobic ion [3, 6].

By voltage-clamp measurements on photolipid-containing planar lipid bilayers (PLBs; folded from 80 m% *E. coli* polar lipid extract (PLE) with 20 m% OptoDArG; schematic of the experimental setup in Figure 2a), we observed an increase in I of the valinomycin–K^+^ complex within milliseconds after the onset of UV light exposure (Figure 2b). Since I measured experimentally is proportional to J of the charged valinomycin–K^+^ complex, I is also proportional to P of K^+^. The ratio I_UV_/I_blue_ was 6.4 *±* 0.8 (mean*±*SEM, N=4), whereby I_UV_ is the current reached following photoisomerization to the photostationary cis state by UV light (average I from 125 ms to 150 ms in Figure 2b) and I_blue_ the current reached following photoisomerization back to the photostationary trans state by blue light (average I from 225 to 250 ms in Figure 2b). In contrast, in the absence of OptoDArG, I_UV_/I_blue_ was merely 1.16 *±* 0.01 (Figure S2a), potentially due to slight changes in membrane temperature (related observation in [8]). The experiment shown in Figure 2b demonstrates that valinomyin–K^+^ transport through photoswitchable PLBs is modulated effectively and reversibly by light.

**Figure 2:**
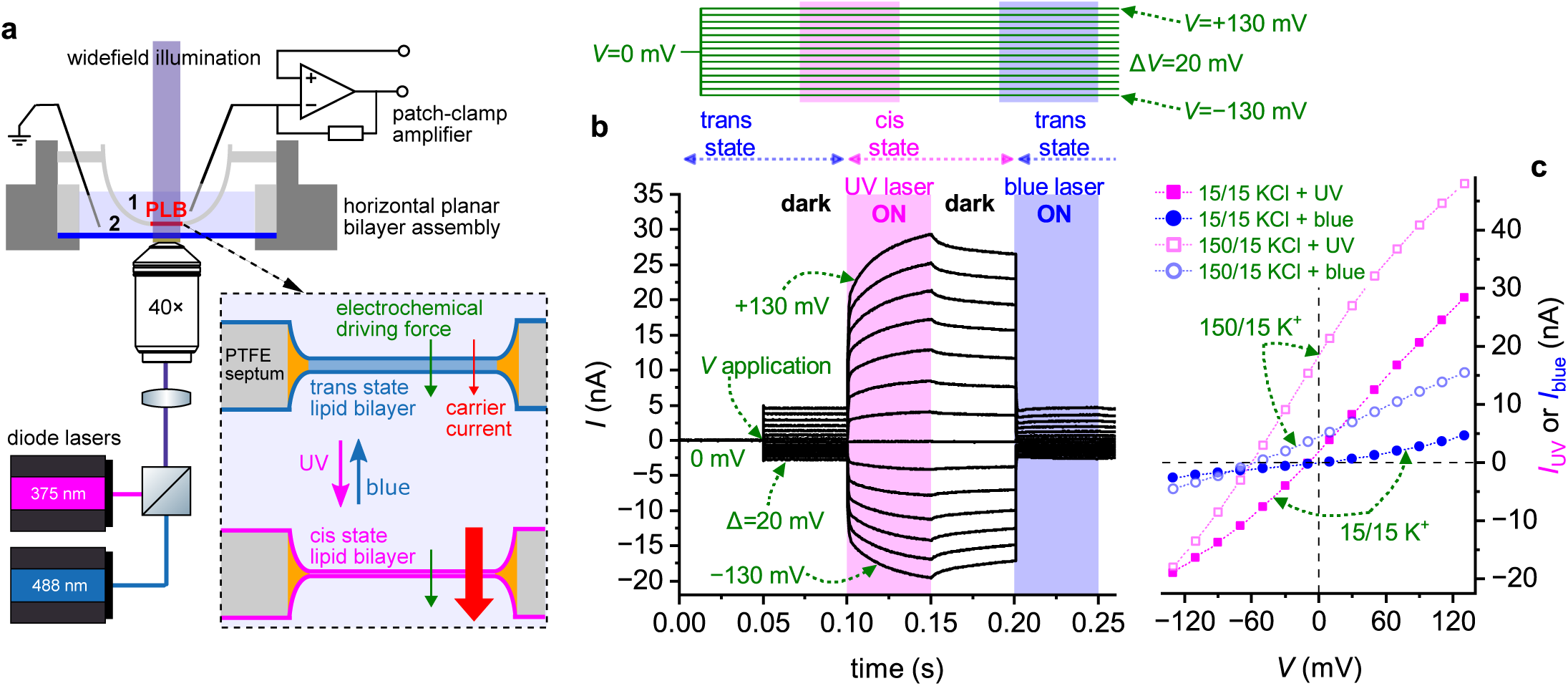
Photoisomerization of photoswitchable PLBs regulates the flux of valinomycin–K^+^ across the membrane. **a,** Schematic of the measurement setup. The patch-clamp amplifier was used to clamp a transmembrane potential, V, across the horizontal PLB and measure the ensuing current, I. A blue and UV laser were used to rapidly isomerize the photoswitchable PLB. The resulting changes in carrier current at constant electrochemical driving force are measured. Side 1 and side 2 is indicated. **b,** Voltage-clamp current, I, recordings on a photoswitchable planar lipid bilayer (PLB) with 10 µM valinomycin added to the aqueous compartments (15 mM KCl, 10 mM HEPES pH 7.4) on both sides. Voltage protocol (inset above): between 0 and 50 ms, V = 0 mV and between 50 and 300 ms, V = 130 mV to *−*130 mV with 20 mV steps between separate sweeps; the delay between consecutively recorded sweeps was 1 s. As indicated, the PLB was exposed to UV light between 100 and 150 ms (magenta bar) and to blue light between 200 and 250 ms (blue bar). UV light evokes a rapid increase in I that is effectively abrogated by blue light exposure. **c,** I–V curves constructed from current traces recorded as in panel **b** by averaging I within certain time intervals and plotting the obtained values over V. I_UV_ values were calculated by averaging I between 125 and 150 ms (magenta squares); I_blue_ values by averaging I between 225 and 250 ms (blue circles). I–V curves are given for symmetric (15 mM KCl at both sides, closed symbols) and asymmetric ionic conditions (150 mM KCl at side 1, 15 mM KCl at side 2, open symbols). The selectivity of I for K^+^ is retained upon UV light exposure.

Current–voltage (I–V) curves constructed from measurements under a 10-fold [K^+^] gradient (Figure 2c) resulted in reversal potentials, V_r_, of *−*56.5 mV (trans state) and *−*59.8 mV (cis state; both determined by interpolation) which is close to *−*59 mV anticipated from the Nernst equation. This shows that the selectivity of valinomycin for K^+^ was retained. Also, we determined that, with variation of the UV laser power (Figure S1b), the fit values for the rate of current increase increase linearly with irradiance (Figure S1c), which is also true for the rate of capacitance change [5]. The increase in I with UV light was clearly the result of photolipid photoisomerization, as evidenced by the rapid abrogation of the current increment by blue light (at 200 ms in Figure 2b; rate ≈ 7000 s*^−^*^1^) which isomerizes the azobenzenes in OptoDArG’s acyl chains back to the trans state. The UV-evoked increase in I does not spontaneously decay to the pre-UV-illumination level without exposure to blue light. This is emphasized by Figure S1a in which UV light was applied but no blue light and I stayed high. Consequently, we can conclude that heating or photodynamic effects did not play a determining role but that changes in bilayer properties that are associated with reversible photolipid photoisomerization did.

Capacitance recordings indicate that the change in bilayer thickness, d, upon photoisomerization did not exceed 10% (Figure S4b). Hence, the dependence on membrane thickness in the traditional one-slab solubility-diffusion model for passive membrane permeation of small molecules – it relates P to the partition coefficient of the substance, K_p_, between the aqueous and the membrane interior, the diffusion coefficient, D_m_, of the substance within the membrane interior, and membrane thickness, d, according to P = K_p_ *·* D_m_/d [9, 1] – cannot account for the observed effects. Likewise, Parsegian’s expression (Equation 1 in [10]) predicts that small changes in d have small effects on P [2, 10]. At <10% change in d, the expected increase in P would be much less than an order of magnitude.

However, we clearly observe much larger current modulation with valinomycin (Figure 2b). A possible explanation for the apparent contradiction between the above predictions and our experimental observations involves alterations in the membrane dipole potential, ϕ_d_. Generated mainly by carbonyl groups of phospholipids and ordered water molecules at the interface, ϕ_d_ (typically around +250 mV inside the membrane) opposes the partitioning of cations and facilitates that of anions [11, 12]. Differences in P between structurally similar organic cations and anions can span up to seven orders of magnitude [11]. A decrease in ϕ_d_ by several tens of millivolts could easily amplify the permeation of the valinomycin–K^+^ complex by an order of magnitude [13]. A drop in ϕ_d_ could result from (i) differences in the dipole moment of the azobenzenes in cis- and trans-OptoDArG (Figure 1e) [14], (ii) a decrease in the density of lipid dipoles due to reduced lipid packing because the cis configuration of the photolipid occupies more space (Figure 1c), and (iii) changes in the orientation of interfacial water or the density of ordered water molecules near the interface [15].

### Regulation of CCCP*^−^* flux across photoswitchable bilayers

To test this hypothesis, we replaced the cationic valinomycin–K^+^ complex with the weak acid carbonyl cyanide *m*-chlorophenylhydrazone (CCCP). The well-known anionic protonophore crosses the membrane in its negatively-charged deprotonated form, CCCP*^−^* [16, 17, 18]. If changes in ϕ_d_ determined the observed light effects, we would expect a decrease in CCCP*^−^* permeability upon exposure to UV light, since lowering ϕ_d_ reduces its favorable effect on anion permeation. However, we observed the opposite: CCCP*^−^* conductance increased with UV illumination (Figure 3a) by a factor I_UV_/I_blue_ = 7.2 *±* 0.1 (mean*±*SEM, N=3), similar to valinomycin–K^+^. Again, H^+^-selectivity was retained, indicated by the retention of a large negative V_r_ in a nominal 2 unit pH gradient (Figure 3b). As before, P was swiftly modulated by light exposure: triggering the blue laser resulted in the immediate abrogation of the UV-induced increment in I owing to a rapid transition to the membrane’s trans state (Figure 3c). These observations exclude changes in ϕ_d_ as the determinant of the approximately one-order-of-magnitude increase in P for valinomycin–K^+^ and CCCP*^−^* following exposure to UV light and the concomitant generation of cis-OptoDArG that thins the membrane (Figure 1c).

**Figure 3:**
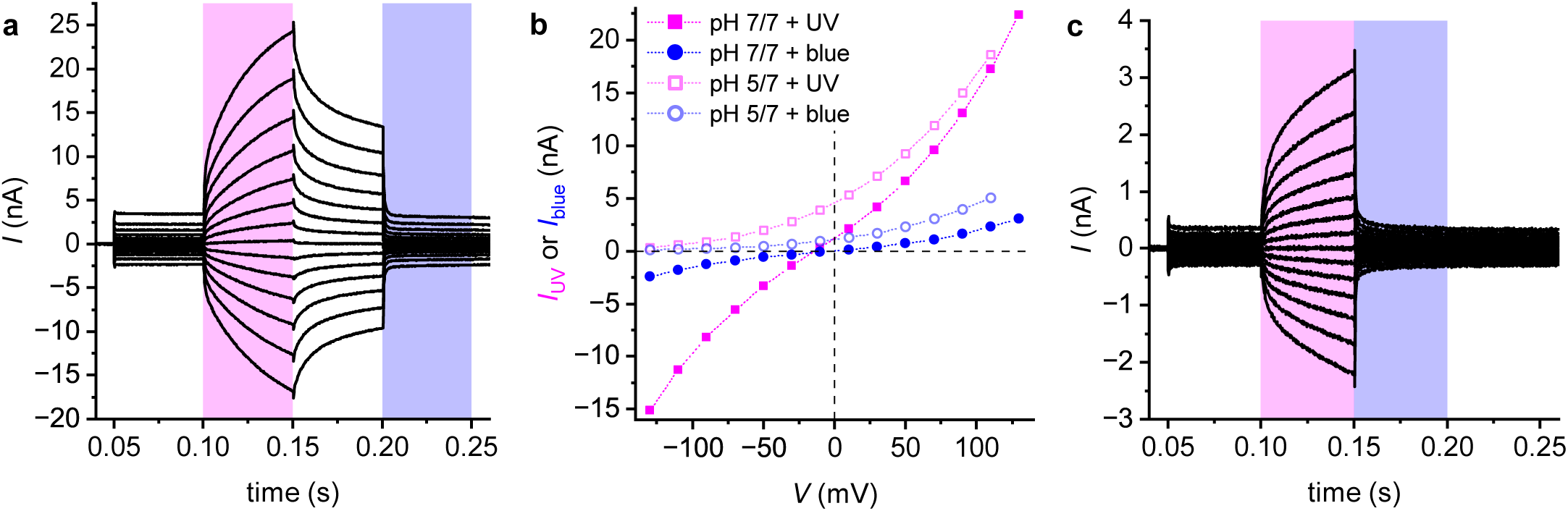
The transmembrane flux of the anionic protonophore CCCP*^−^* is rapidly modulated by photoisomerization of membrane-embedded photolipids. **a,** Voltage-clamp current recordings on a photoswitchable PLB with 10 µM CCCP added to the aqueous compartments (100 mM KCl, 20 mM HEPES pH 7.0) on both sides. The recordings were made as in Figure 2b. As with valinomycin–K^+^, I increases upon UV light exposure. That is, the flux of CCCP*^−^* is increased across the cis state membrane. **b,** I–V curves constructed as described in Figure 2c for recordings under symmetric (pH 7 at both sides; closed symbols) and asymmetric conditions (pH 5 at side 1, pH 7 at side 2; open symbols). Upon creating the gradient in pH by the addition of HCl to compartment 1, V_r_ shifted towards large negative values, consistent with proton-selective transport. Whilst UV light exposure increased I, the current remained selective for H^+^. **c,** Photoswitchable PLB with 2.5 µM CCCP added to the aqueous compartments (15 mM KCl, 10 mM HEPES pH 7.4) on both sides. Voltage protocol as in panel **a**. In contrast to panel **a**, UV light exposure between 100 and 150 ms was immediately followed by blue light from 150 to 200 ms. This record shows that blue light leads to the immediate abrogation of the UV-evoked increment in I as a result of switching to the trans bilayer state.

Hence, we observed ≈7-fold amplifications in the flux of valinomycin–K^+^ and CCCP*^−^* in the cis bilayer state. That is, the effect appears to be independent of carrier charge. To that effect, returning to Parsegian [2] and Dilger, McLaughlin, McIntosh, and Simon [10], we speculated that changes in the dielectric constant of the bilayer interior, ɛ_hc_, could be held accountable. In fact, even a seemingly modest increase in ɛ_hc_ from 2 to ≈4.5 led to a >1000-fold increase in the conductance, g, of thiocyanate but had a much smaller effect (10-fold increase in conductance) on nonactin–K^+^, another cationic K^+^ ionophore [10]. Consequently, the results on nonactin in [10] suggest that ɛ_hc_ would have to increase pronouncedly to account for the ≈8-fold increase in g that we observe with valinomycin. However, ɛ_hc_ also appears in the expression for the electrical capacitance of the PLB, C_m_ = ɛ_0_ ɛ_hc_ A_m_/d, where ɛ_0_ is vacuum permittivity and A_m_ bilayer surface area. We did not observe a corresponding disproportionately large increase in C_m_ upon transitioning to the bilayer cis state (Figure S4b), which argues against large changes in ɛ_hc_.

We tested the hypothesis regarding changes in ɛ_hc_ by assessing the transport of ions across our photoswitchable PLBs. First, we determined the effect of OptoDArG photoisomerization on background ion conductance and found that g increases from 12.1 *±* 0.8 pS to 20.7 *±* 0.8 pS in the cis bilayer state (Figure 4a). The corresponding increase in 71% is notable, but by an order of magnitude lower than the effect on valinomycin–K^+^ and CCCP*^−^*. Second, we measured transport of tetraphenylborate anions (TPB*^−^*) across trans and cis state bilayers. TPB*^−^* is a hydrophobic ion that permeates the membrane in its anionic form [19, 12]. P of TPB*^−^* is much higher than that of inorganic ions, and the transport of TPB*^−^* towards the membrane limits the steady-state current. At the TPB*^−^* concentrations used (200 nM), application of a voltage (V = 120 mV at t = 0 ms in Figure 4c) leads to the redistribution of membrane-adsorbed TPB*^−^* between the adsorption sites in each leaflet (schematic in Figure 4b), resulting in an exponentially decaying current [19]. A monoexponential fit to the transient currents in Figure 4c allowed us to estimate the initial current, I_0_, and time constant of the transient, τ [20]. Both parameters increase by a factor of ≈2 in the cis state membrane (inset in Figure 4c). This indicates that the rate of transport between the adsorption sites increases in the presence of cis-OptoDArG, but the increase is small compared to the effects on valinomycin–K^+^ and CCCP*^−^* currents. From these observations, we conclude that transport rate amplifications may not be the basis for the effect of photoisomerization on valinomycin–K^+^ and CCCP*^−^*.

**Figure 4:**
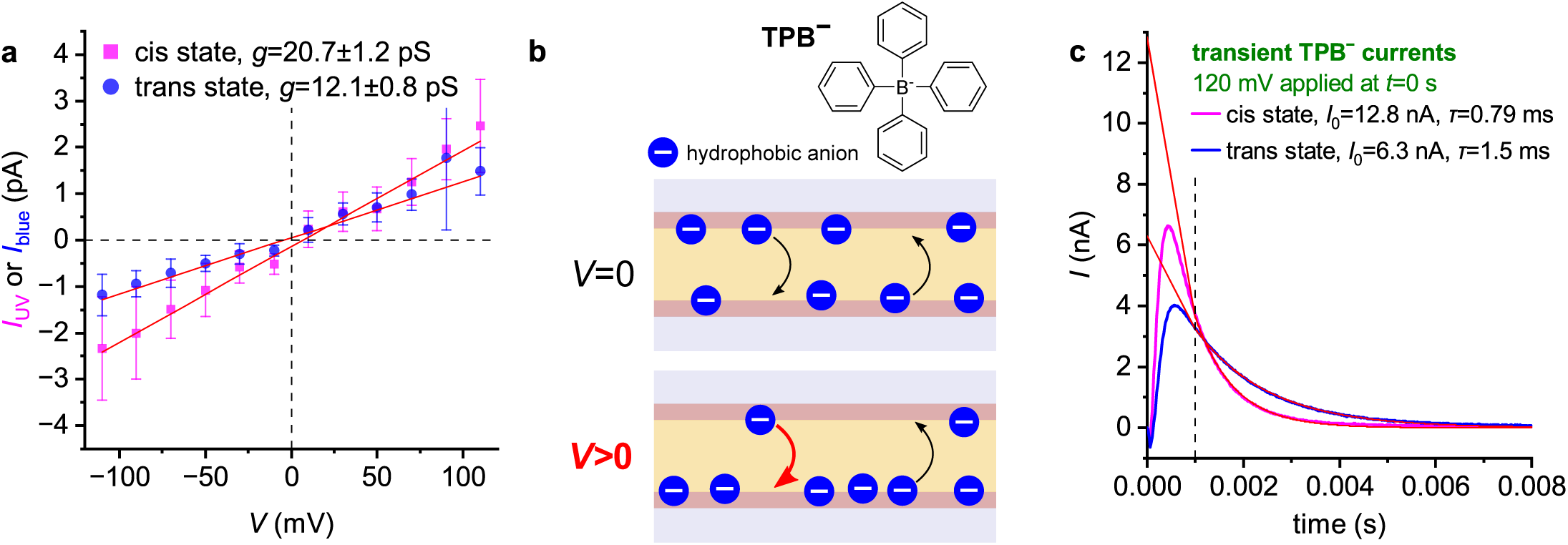
Photoisomerization has a comparatively small effect on background and hydrophobic ion conductance. **a,** To infer the change in background ion conductance upon photoisomerization, recordings as in Figure 2b but in the absence of carrier were conducted; the buffer was 15 mM KCl, 10 mM HEPES pH 7.4. I–V curves were constructed from the obtained records, as described in Figure 2c, and data points from 5 separately prepared experiments were averaged (error bars correspond to SD). The curves were fit by linear models with offset, with each point weighted by 1/SD^2^ (R^2^ of 0.97 (cis state) and 0.96 (trans state)). Background ion conductance increases by ≈71% in the bilayer cis state, a small effect compared to the experiments with valinomycin and CCCP. **b,** Tetraphenylborate (TPB*^−^*) is a hydrophobic anion that permeates the membrane. At low bulk TPB*^−^* concentrations, voltage application leads to the redistribution of membrane-adsorbed ions between the leaflets without considerable contribution from the bulk [19]. Hence, in contrast to carriers, chemical uptake and release reactions at the interface play little role (cf. Figure 1a). **c,** Transient TPB*^−^* currents evoked upon the application of V = 120 mV at t = 0 s with the bilayer in the cis (magenta lines) or trans state (blue lines); TPB*^−^* was 200 nM in 100 mM NaCl, 10 mM HEPES pH 7.4 (conditions similar to ref. [19]). Each transient was recorded twice with a delay of 800 ms to check for steady-state conditions: the curves overlap perfectly. Due to a 2 MΩ resistance in series with the PLB, necessary for capacitance compensation, and the deployed 10 kHz Bessel filter, the transients are smoothed. Hence, to estimate initial current I_0_ and decay time τ, data points from 1 to 8 ms were fit with a monoexponential model. The fits (red lines) indicate a modest effect (around 2-fold) of bilayer state on I_0_ and τ, which is small compared to the experiments with valinomycin and CCCP.

To reconcile the surprising finding of differential effects of photoisomerization on currents mediated by carriers and ions, we looked at where they differ. Although both permeate the membrane in a charged state, carriers additionally undergo association and dissociation reactions with their respective target ions at the interface – the corresponding rates are k_A_ and k_D_. Hence, one may imagine that photoisomerization affects these interfacial uptake and release reactions. Qualitatively, the idea here is similar to the departure from the traditional one-slab solubility-diffusion model, which assumes partitioning into a homogeneous hydrophobic membrane, to a three-slab model [1]. In this model, the membrane is divided into two headgroup regions and a hydrophobic core between, each offering a different resistance to permeation. The total permeability of the membrane, P, is determined by the slab with the highest resistance. Within narrow confines, Nagle, Mathai, Zeidel, and Tristram-Nagle [21] could explain the permeation of water by such a model: lipid unsaturation increases headgroup spacing, allowing water molecules to traverse the headgroup region and reach the hydrophobic core more easily. As a consequence, the water permeability of membranes composed of unsaturated lipids can be up to an order of magnitude higher than that of membranes with a more saturated composition [22]. Importantly, a similar effect has been observed with valinomycin, where k_A_ increased and k_D_ decreased with increasing lipid unsaturation [23]. Faster uptake and slower release of K^+^ – reflected in an increase in the ratio k_A_/k_D_ from 1.5 (monoglycerides with oleoyl (C_18:1_) chains) to 9 (linolenoyl (C_18:3_) chains) [23] – may increase the number of membrane-adsorbed valinomycin–K^+^ and transmembrane flux. In fact, a factor of 9 is similar to the increase in valinomycin–K^+^ current through cis state PLBs observed here, lending credibility to the hypothesis.

That is, our model is the following (Figure 5): the switch from trans to cis configuration of membrane-embedded OptoDArG mimics structural changes due to increasing lipid unsaturation. Like unsaturated lipids, cis-OptoDArG occupies more space laterally (Figure 1b,c), increases headgroup spacing, and thus conceivably decreases the resistance posed by the headgroup region. In the case of valinomycin, this may facilitate the access of K^+^ to interfacially-adsorbed carriers, accelerating interfacial uptake and release reactions. A similar mechanism may apply to CCCP, whereby both the access of hydronium ions to interfacially-adsorbed carriers and the partitioning of CCCP into the bilayer may be enhanced, increasing overall permeability. In fact, the slower UV-induced increase in I with CCCP (Figure 3a) compared to valinomycin may indicate just that: a gradual increase in the intramembrane concentration of CCCP upon exposure to UV light by partitioning of CCCP from the bulk into the more permissive cis state membrane.

**Figure 5:**
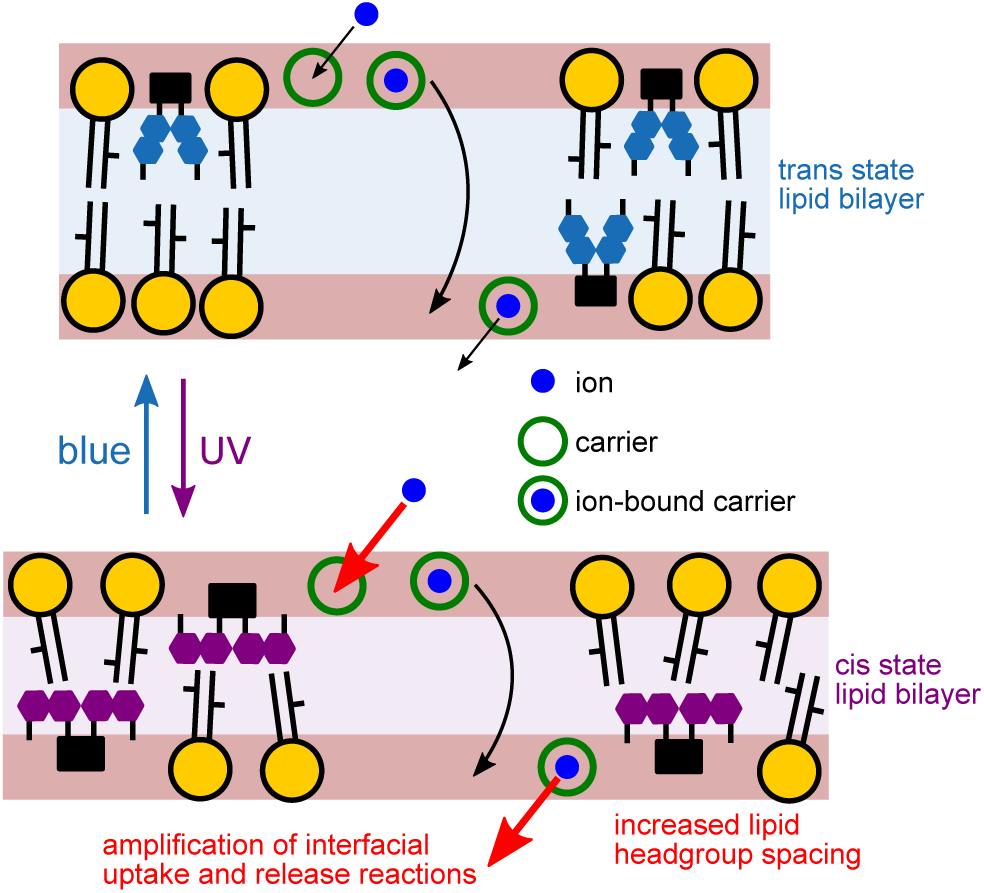
*Model:* Regulation of carrier currents by light-driven modulation of lipid packing, interfacial reactions and partitioning. Our results on carrier and (hydrophobic) ion transport reported here are consistent with the view that photoisomerization of membrane-embedded photolipids alters lipid packing and thereby affects interfacial uptake and release reactions between membrane-adsorbed carrier molecules and ions in solution. This could, for example, result in an increase in k_A_/k_D_ and therefore the number of membrane-adsorbed ion-bound carrier molecules, increasing the overall flux. Because of the relative excess of uncharged carrier molecules at the interface, amplification of the rate of ion-bound carrier transport between the adsorption sites is not necessary. This model agrees qualitatively with a three-slab partitioning scheme where photolipid photoisomerization alters partitioning at the level of the headgroup regions but has little effect on the transport rate across the hydrophobic core.

### NB-lipid proton flux is effectively modulated by bilayer photoisomerization

In an attempt to maximize the light effect, we sought to reduce the dark current (I when the photolipid is in its trans state). To explore this, we replaced the anionic protonophore CCCP with a cationic protonophore, a weak base that permeates the membrane in its protonated positively-charged state. The rationale is that ϕ_d_ facilitates the movement of anionic protonophores across the membrane while opposing that of cationic protonophores, thus reducing the corresponding dark current. We selected Nile Blue (NB), a benzophenoxazine-based red fluorescent dye, which has been described as a mitochondrial uncoupler [24], but is relatively inefficient as a protonophore. To make NB more hydrophobic and thus create a situation similar to that with valinomycin, we attached an acyl chain to Nile Blue, creating NB-lipid (Figure 1d). Upon the addition of NB-lipid, to the aqueous solutions on both sides of the membrane, a gradual increase in membrane conductance, g_0_, was observed (Figure 6a,b). The recorded I–V curves were characteristically supralinear, which is the result of ion permeation across the trapezoidal potential barrier ϕ_d_ [1]. Hence, NB-lipid indeed showed protonophoric activity and gradually incorporated into the lipid bilayer from its aqueous surroundings.

**Figure 6:**
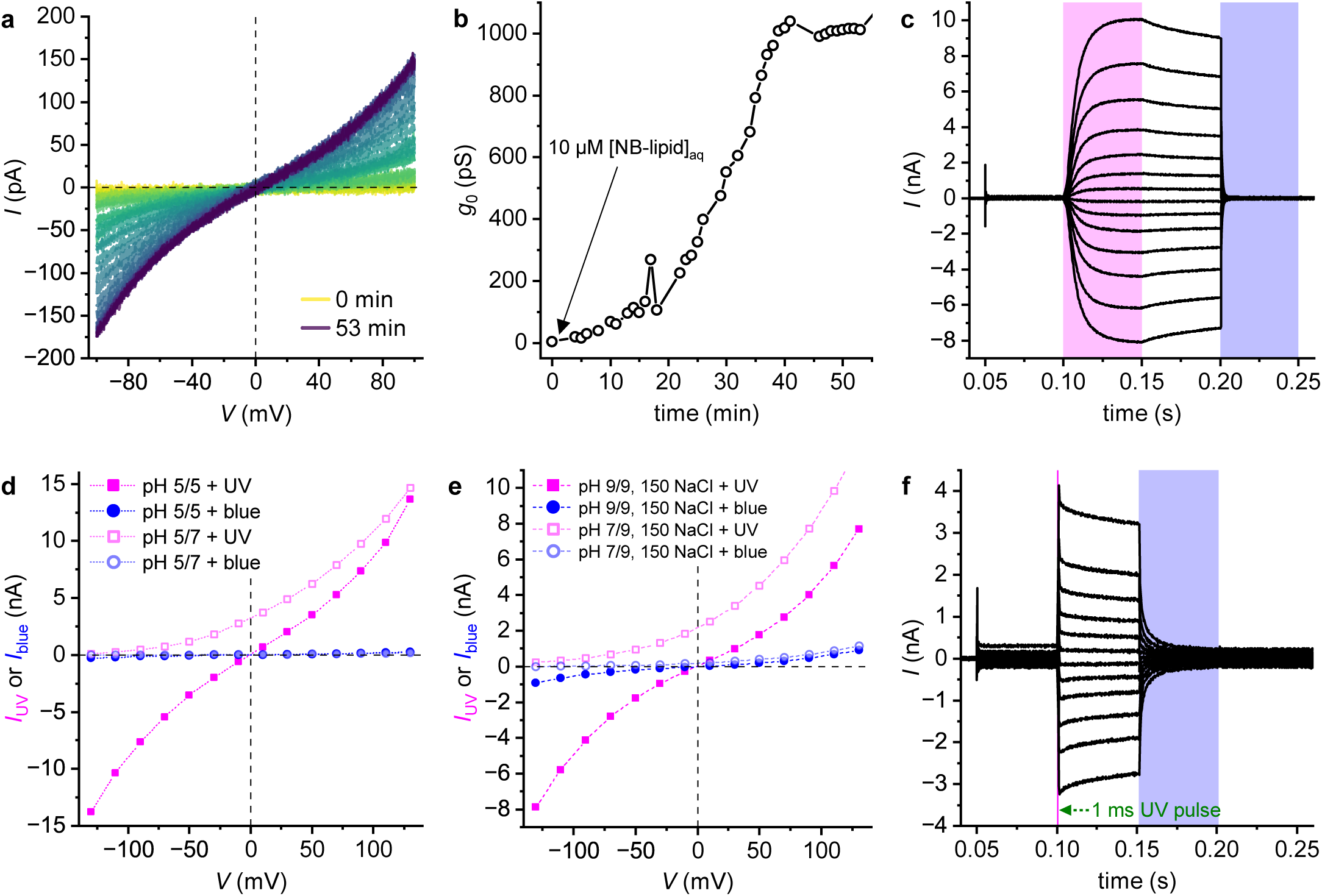
NB-lipid incorporates into lipid bilayers and can be regulated very efficiently by photolipid photoisomerization. **a,** Voltage ramps measured on a vertical 135 µm-diameter photoswitchable PLB folded from 80 m% *E. coli* PLE and 20 m% OptoDArG in 15 mM KCl, 10 mM HEPES pH 7.4. 10 µM [NB-lipid]_aq_ was added to both compartments (stirred continuously) immediately after the record at t = 0 min (yellow line) and further voltage ramps were recorded under continuous stirring for 53 min. Voltage protocol: 0.5 s at *−*100 mV, increase to 100 mV at 100 mV s*^−^*^1^, 0.5 s at 100 mV; the constant-voltage segments are not shown. **b,** The cubic I–V curves in panel **a** were fit by the following equation: I(V) = g_0_ *·* (1 + α V ^2^) *·* V + o; where I(V) is the current at a particular voltage, g_0_ the conductance at V = 0 mV and α is a supralinearity factor [1]; o accounts for a small current offset. The obtained g_0_ values are plot over time of record. Conductance increased gradually, indicating membrane incorporation of aqueous NB-lipid. **c,** Voltage-clamp current recordings on a photoswitchable PLB containing 1 m% NB-lipid in a solution containing 15 mM KCl, 10 mM HEPES pH 7.4. Recordings were made as in Figure 2b. In this record, exceptionally high on–off current regulation by light (>200-fold) was achieved. **d,** I–V curves constructed from current records as described in Figure 2c. Conditions were symmetric (100 mM KCl, 20 mM MES, pH 5 at both sides; closed symbols) and asymmetric (pH at side 2 increased to 7; open symbols). NB-lipid currents are proton-selective. **e,** Even at physiological salt concentrations (150 mM NaCl, 20 mM TRIS, pH 9), H^+^-selectivity is retained. This can be appreciated from the large negative-going shift in V_r_ upon reducing pH in compartment 1 by the addition of acid. **f,** Recordings made as in panel **c** whereby the duration of UV light exposure was reduced to 1 ms (V ranged from 110 mV to *−*110 mV). Even short pulses of UV light can effectively regulate membrane proton permeability.

At lower concentrations of NB-lipid (established by adding 1 m% NB-lipid to the PLB-forming lipid mixture), g_0_ in the presence of trans-OptoDArG could also be much lower. For the record shown in Figure 6c, a cubic fit of I_blue_ over V resulted in g_0_ ≈150 pS. Critically, upon UV illumination, we observed a >200-fold increase in I and hence H^+^ permeability. Although replicates of this experiment (freshly prepared measurement chambers, different lipid mixtures; Figure S4a) revealed pronounced variations in g_0_ (4.8*±*2.1 nS (mean*±*SEM, N=5)) and the fractional change in I upon photoisomerization (I_UV_/I_blue_=70*±*40 (mean*±*SEM, N=5)), dark current was generally small and I_UV_/I_blue_ large.

Experiments carried out with a transmembrane proton gradient confirmed that the current is proton-selective (Figure 6d). Furthermore, this selectivity remained intact even in the presence of physiological salt concentrations (150 mM NaCl in Figure 6e; 150 mM KCl in Figure S3). Finally, even short UV light pulses (1 ms long) resulted in a sizable increment in I (Figure 6f); this may be important for cell applications as it reduces the amount of energy delivered. What may be the reason for the enhanced sensitivity to photolipid state of NB-lipid compared to valinomycin and CCCP? We noted that NB-lipid resembles a modified fatty acid. Hence, we imagined that looser lipid packing may amplify both interfacial proton uptake and release, according to our model (Figure 5), as well as the rate of flip-flop between the bilayer leaflets.

Notably, the combination of NB lipid and photolipids partially outperforms ChRs – an alternative method to generate light-switchable proton currents [25, 26, 27, 28, 29]. At neutral pH, the photocurrents generated are comparable in magnitude (Figure 6c) to those elicited by the human proton channel H_V_1 or ChR in transfected cells [30, 31]. As in the case of H_V_1 or ChR2, the current is highly selective for protons. However, the ratio of proton to sodium permeability, P_H_^+^ /P_Na_^+^, for ChR2 was only 2 *×* 10^6^ to 6 *×* 10^6^, resulting in a dominant Na^+^ component at neutral pH in mammalian brains [29]. In contrast, even in a high-salt environment, the contribution of the salt to the reversal potential of a NB-lipid-containing cis state membrane was negligible (Figure 6e).

## Conclusion

The photolipid-based approach at regulating carrier currents valuably distinguishes itself from (i) photopharmacological approaches that aim to regulate drug activity by light [32, 33] and which typically focus on classical orthosteric or allosteric ligands to gate protein activity (e.g., ion channels and other membrane transporters) [34, 35], (ii) “molecular machines for controlling transmembrane ion transport” [36] where light-dependent currents are generated and ionophores *per se* are modified with light-sensitive moieties, (iii) rotary molecular motors which have been reported to increase membrane ion permeability upon irradiation [37, 38], but have been shown to act at least partially through irreversible photodynamic membrane lipid peroxidation [39] rather than drilling [40], (iv) artificial supramolecular channels [41, 42], and (v) light-based approaches at regulating mechanosensitive channels by alterations in bilayer mechanical properties [4, 43, 5, 44]. Our approach is all-chemical and allows for light-dependent, millisecond-scale regulation of well-established highly-selective carriers that have been in use for decades. It achieves large currents, a submillisecond photoresponse with long-lasting effect, easily tuned ion scope, and the potential to rationally shift its single photon action spectra from the UV/Vis right up to the NIR/SWIR [45, 35, 46].

Unlike the ChR-based optogenetic approach [25, 26, 27, 28, 29], which requires genetic transfection to achieve selective ion currents in response to incident light, our purely chemical approach avoids this complication since both NB-lipid and photolipids [5, 47] can be administered via aqueous solution. We believe that our approach can complement optogenetics and that the tailored design of other carriers may eventually allow efficient regulation of Na^+^ and K^+^ permeabilities with light without the need for genetic transfection, much like the pursuit for optimizing ChR function [28, 29]. We imagine photolipid-regulated carriers as a complementary and convenient all-chemical approach toward achieving light-based ion permeability regulation in the biological and synthetic cell or the artificial neuron.

In summary, we demonstrated that photoisomerization of photoswitchable bilayers, which reduces their thickness, can result in significant permeability alterations of up to two orders of magnitude. Our results are consistent with modulation of interfacial uptake and release reactions between carriers and their respective cargoes because of altered lipid packing upon photoisomerization. Qualitatively, this resembles a three-slab model in which the lipid headgroup regions drive the effect, while the inner core behaves as predicted by Parsegian [2]. This reveals that the apparent contradiction with his original analysis initially attested is only superficial.

## Acknowledgments

This research was funded in whole by the Austrian Science Fund (FWF) grants P36399 and P34826 to Peter Pohl.

## Supplementary Information

### S1 Material and Methods

#### S1.1 Materials for planar lipid bilayer experiments

*E. coli* Polar Lipid Extract (PLE, item no. 100600) was obtained from Avanti Polar Lipids (distributed by Merck) and kept at *−*80 °C. OptoDArG was synthesized by the group of Dr. Glasnov as described previously [1]. Lipid aliquots and mixtures were prepared within amber glass micro reaction vials from lipids dissolved in chloroform. Prior to storage at *−*80 °C, solvent was evaporated by a mild vacuum gradient (Rotavapor, Büchi Labortechnik AG) and the dried lipids were flooded with argon. All aqueous buffers used in the PLB experiments were freshly prepared from laboratory-grade dry substances (supplied by VWR, Merck, or Fisher Scientific) dissolved in ultrapure water (> 18 MΩ cm, Milli-Q water purification system) and pH-adjusted using a daily-calibrated pH meter (FiveEasy, Mettler Toledo). CCCP (carbonyl cyanide *m*-chlorophenylhydrazone, 98%, Thermo Scientific) and valinomycin were kept as stocks in DMSO. Sodium tetraphenylborate (>99.5%, T25402, Sigma-Aldrich) was added from a stock solution in ethanol. NB-lipid was synthesized in a parallel study, with synthesis and characterization given there as molecule **S43** [2]. NB-lipid was kept in a dried state at *−*80 °C and aliquots were prepared in DMSO prior to measurement.

#### S1.2 Horizontal planar lipid bilayer experiments

Planar lipid bilayer (PLB) experiments with laser irradiation for rapid photoisomerization were conducted as recently described [3]. A schematic of the setup is shown in Figure 2a. Solvent-depleted horizontal PLBs (specific capacitance 0.75 µF cm*^−^*^2^) were folded from lipids spread on top of aqueous buffer in the lower and upper compartment of a custom-made chamber assembly made of PTFE [4, 5]. One of the following lipid mixtures were used in this study: photoswitchable PLBs: 80 m% *E. coli* PLE and 20 m% OptoDArG; photoswitchable PLBs with 1 m% NB-lipid: 79 m% *E. coli* PLE, 1 m% NB-lipid and 20 m% OptoDArG; regular PLBs: 100 m% *E. coli* PLE; NB-lipid-containing PLBs: 99 m% *E. coli* PLE and 1 m% NB-lipid.

First, an aperture of around 70 µm in diameter in 25 µm-thick PTFE foil (Goodfellow GmbH) was created by high voltage discharge – this prepares the septum separating the macroscopic compartments. In this study, diameters were between 70 µm to 85 µm, with exceptions denoted explicitly. The PLB diameters given in the main text refer to the size of this aperture. After the septum was treated with 0.6 vol% hexadecane in hexane, hexane was allowed to evaporate for >1 h. The residual hexadecane facilitates the solvent annulus or torus that later laterally anchors the PLB within the aperture [6]. The septum was attached by silicon paste to the lower side of the upper compartment of the chamber assembly. Lipids at the air–water interfaces were prepared by applying lipid mixtures dissolved in hexane at a concentration of 10 mg mL*^−^*^1^ onto both aqueous interfaces. After hexane had evaporated, a horizontal PLB was folded by rotation of the upper compartment of the chamber assembly.

A 30 mm-diameter cover glass (No. 1, Assistent, Hecht Glaswarenfabrik GmbH & Co KG) fixed with a threaded PTFE ring comprised the bottom of the lower compartment. The chamber assembly was installed on the sample stage of an Olympus IX83 inverted microscope equipped with an iXon 897 E EMCCD (Andor, Oxford Instruments Group). The chamber holder was equipped with screws for fine translation of the upper compartment in z direction to position the horizontal PLB within the working distance of a 40×/1.30 NA infinity-corrected plan fluorite oil immersion objective (UPLFLN40XO, Olympus) or a 40×/0.65 NA infinity-corrected plan achromat air objective (PLN40X, Olympus). The motorized microscope and real-time controller (U-RTC, Olympus) used for synchronizing lasers and electrophysiological acquisition were controlled using the proprietary cellSens software (Olympus).

For electrical measurements, a Ag/AgCl agar salt bridge containing 0.5 M KCl was put into each compartment and connected to the headstage of an EPC 9 patch-clamp amplifier (HEKA Elektronik, Harvard Bioscience). Headstage and chamber assembly were housed in a Faraday cage. Voltage-clamp measurements were conducted using PATCHMASTER 2x91 software (HEKA Elektronik, Harvard Biosciences). Current was analogously filtered at 10 kHz by a combination of Bessel filters and acquired at 50 kHz. Amplifier offsets were corrected by subtracting the average current recorded at V = 0 mV under symmetric conditions. Data recorded with PATCHMASTER was exported, analyzed, and graphed using Mathematica 14 (Wolfram Research) and OriginPro 2024 (OriginLab Corporation).

Rapid photoisomerization of photolipids embedded in horizontal PLBs was achieved by exposure to blue (488 nm diode laser, iBEAM-SMART-488-S-HP, TOPTICA Photonics) and UV laser light (375 nm diode laser, iBEAM-SMART-375-S, TOPTICA Photonics). Both lasers were digitally modulated and separately focused into the back-focal plane of the objective via the ZT488/640rpc main dichroic mirror (Chroma). The diameter of the blue laser profile at the sample stage was ≈58 µm (1/e^2^) whilst the UV laser profile spanned roughly 150–200 µm (its shape was less defined owing to the absence of a spatial filter). At a software-set output power of 200 mW (blue) and 70 mW (UV), ≈20–30 mW (blue and UV) exited the microscope objective depending on current alignment, determined by a photodiode (S120VC, Thorlabs).

#### S1.3 Vertical planar lipid bilayer experiments

Vertical PLBs were folded in a similar manner to the description above and as given in [7]. The Faraday cage was equipped with a magnetic stirrer positioned underneath the PTFE chamber, which allowed the aqueous compartments on both sides of the vertical PLB to be stirred by magnetic PTFE-coated stir bars. Ag/AgCl agar salt bridges were used. The recordings were made with an EPC 10 patch-clamp amplifier (HEKA Elektronik, Harvard Bioscience).

#### S1.4 Biexponential function fits

To extract rate information from the time course of current upon light exposure, we fitted several sets of curves with the following bi-exponential equation:

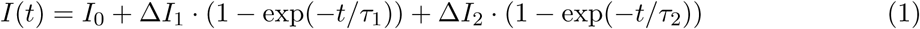

where I_0_ denotes basal current, ΔI_1_ and ΔI_2_ the increment in I associated with growth at rate 1/τ_1_ and 1/τ_2_. Typically, several curves were fit globally (Mathematica resource function MultiNonlinearModelFit) and curves recorded at lower voltages (*|*V *| ≤* 30 mV) were neglected.

### S2 Supplementary Figures

**Figure S1:**
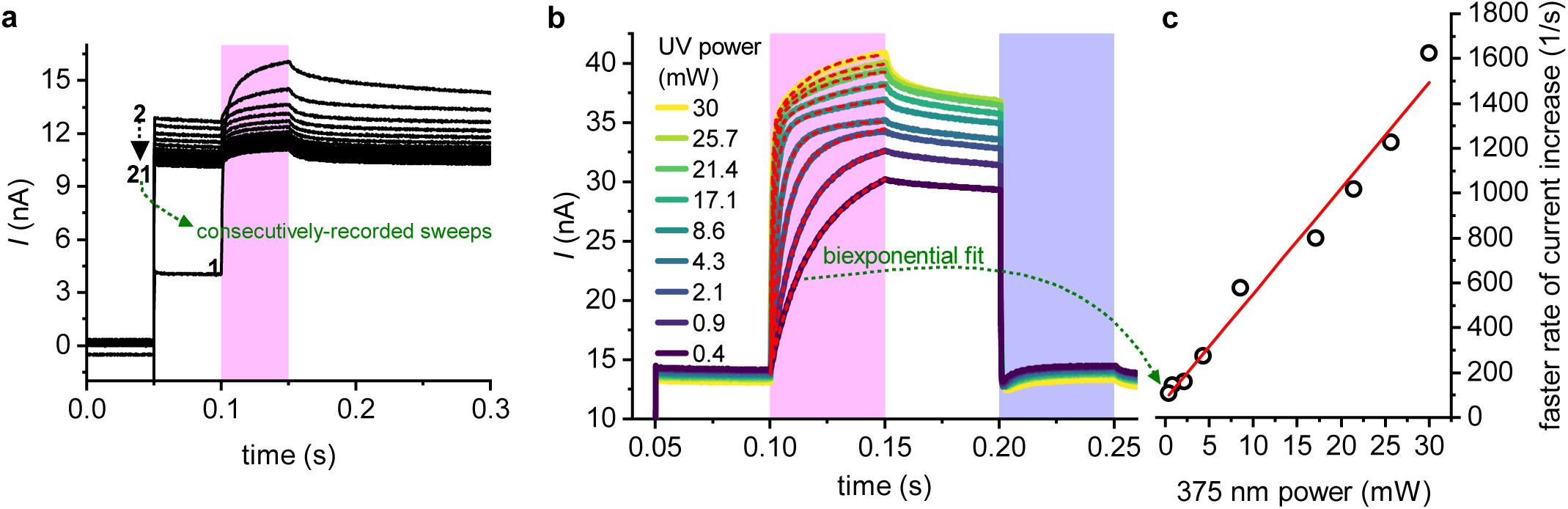
**a,** Current recordings on a photoswitchable PLB with 10 µM valinomycin on both sides (15 mM KCl, 10 mM HEPES pH 7.4). The following was repeated 21 times with an intersweep delay of 1 s: between 0 and 50 ms, V = 0 mV and between 50 and 300 ms, V = 130 mV; between 100 and 150 ms the PLB was exposed to UV light (no blue light exposure). The UV-evoked current increment does not decay spontaneously without blue light because cis-azobenzene does not decay at this timescale [8]. **b,** Irradiance-dependence of the UV light-triggered increment in I with valinomycin. Photoswitchable PLB with 10 µM valinomycin in 150/15 mM KCl, 10 mM HEPES pH 7.4. Recordings were conducted as in panel Figure 2b but with V constant at +130 mV whilst the power of the UV laser was varied in consecutive sweeps; the given values are power at the sample stage. The red-dashed lines are biexponential fits. I increases at a rate proportional to irradiance. **c,** Plot of the faster rate of change of I fitted in panel **b** over respective incident UV laser power. The red line is a fit to a linear model with offset (R^2^ = 0.98).

**Figure S2:**
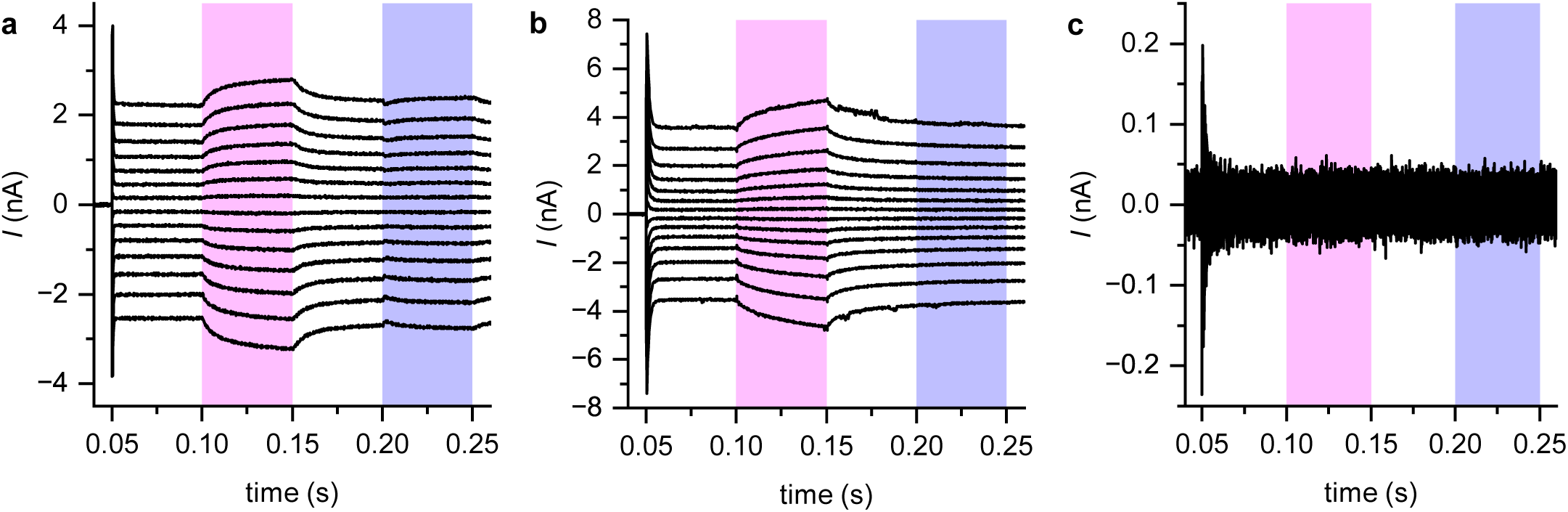
Voltage-clamp current recordings on non-photoswitchable PLBs folded from *E. coli* PLE bathed in 15 mM KCl, 10 mM HEPES pH 7.4 containing nominally 10 µM valinomycin (panel **a**) or 10 µM CCCP (panel **b**). The records in panel **c** were made on a membrane folded from 99 m% *E. coli* PLE with 1 m% NB-lipid. Recordings were made as in Figure 2b.

**Figure S3:**
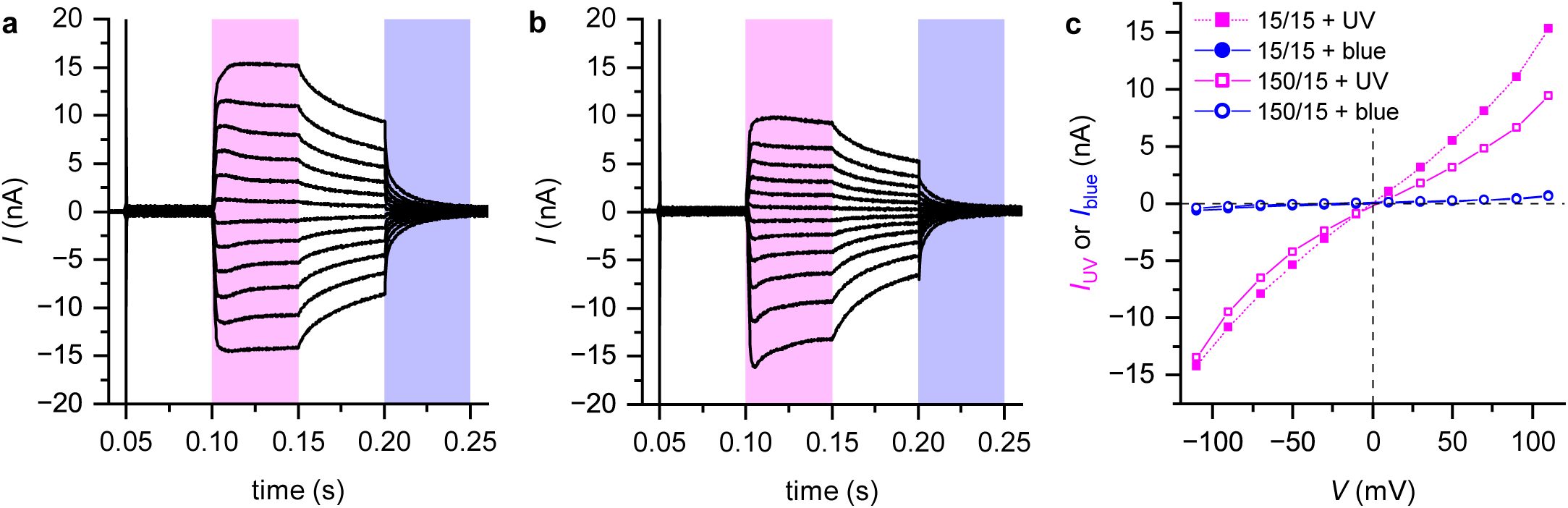
**a,** Voltage-clamp current recordings on a 85 µm-diameter photoswitchable PLB folded from 79 m% *E. coli* PLE, 20 m% OptoDArG and 1 m% NB-lipid in 15 mM KCl, 10 mM HEPES pH 7.4. Recordings were made as in Figure 2b with V ranging from 110 mV to *−*110 mV. **b,** K^+^ concentration on side 1 was made 150 mM by the addition of 3 M KCl solution. Recordings were made as in panel **a**. **c,** I–V curves constructed from different intervals of the current traces in panel **a** and **b**, as described in Figure 2c.

**Figure S4:**
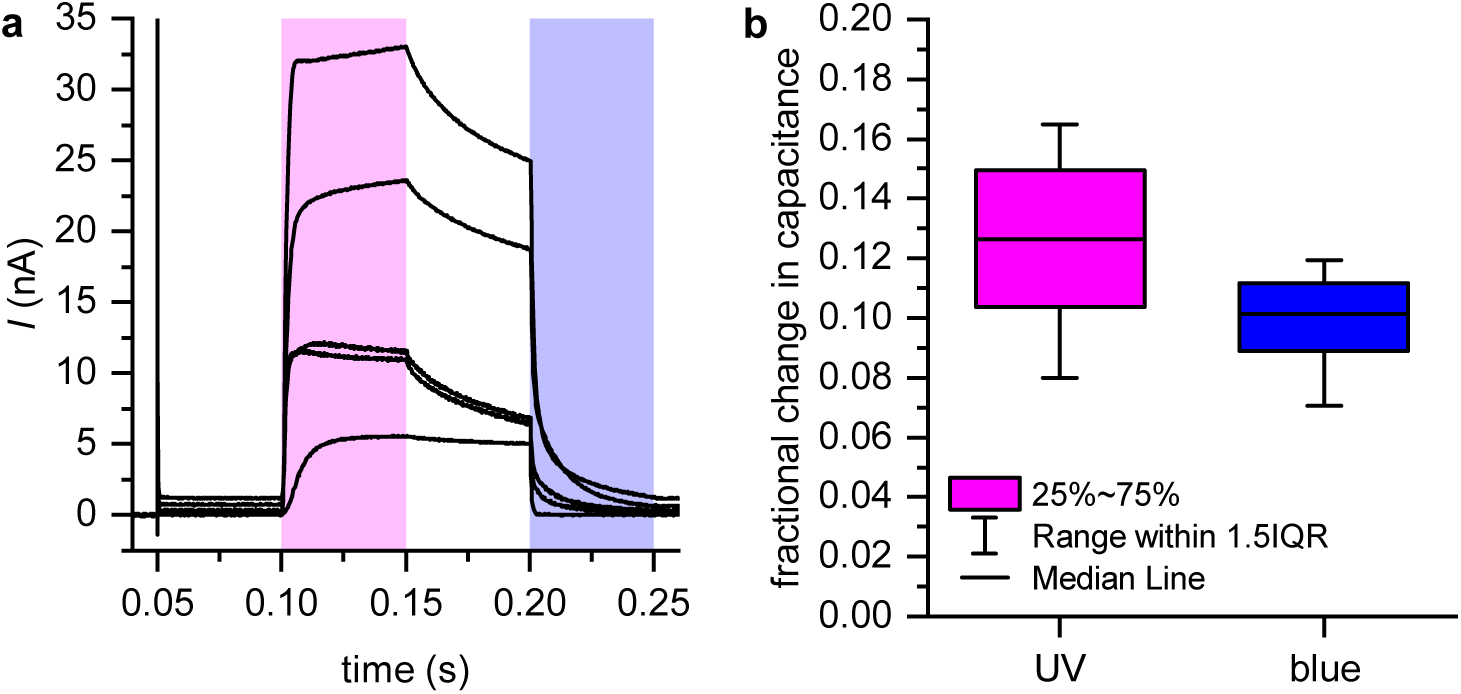
**a,** Voltage-clamp current recordings on 75 to 95 µm-diameter photoswitchable PLBs folded from 79 m% *E. coli* PLE, 20 m% OptoDArG and 1 m% NB-lipid in 15 mM KCl, 10 mM HEPES pH 7.4. The traces were recorded in separate measurements which were equally conducted. V = 90 mV was applied at 50 ms. Light exposure as in Figure 2b. **b,** Fractional changes in bilayer capacitance, C_m_, upon UV and blue light exposure. PLBs were folded from 80 m% *E. coli* PLE + 20 m% OptoDArG (11 recordings from 3 separate experiments). Capacitance recordings were conducted as described in Bassetto, Pfeffermann, Yadav, Strassgschwandtner, Glasnov, Bezanilla, and Pohl [3] (Figure 2a,b there). Fractional change in capacitance was calculated as 1 *−* C_m,peak_/C_0_ whereby C_m,peak_ denotes peak capacitance achieved within milliseconds after light exposure and C_0_ capacitance immediately before light exposure.

